# Dynamics of primary productivity in relation to submerged vegetation of a shallow, eutrophic lagoon: a field and mesocosm study

**DOI:** 10.1101/2021.02.12.430911

**Authors:** Maximilian Berthold, Martin Paar

## Abstract

Aquatic ecosystems nowadays are under constant pressure, either from recent or historical events. In most systems with increased nutrient supply, submerged macrophytes got replaced by another stable state, dominated by phytoplankton as main primary producer. Yet, reducing the nutrient supply did not yield the aimed goal of restored habitats for submerged macrophytes in systems worldwide. The question arises, why submerged macrophytes do not re-colonize, and if they are actually competitive. Therefore, primary production assays were conducted in *ex-situ* bentho-pelagic mesocosms and compared to the actual ecosystem, a turbid brackish lagoon of the southern Baltic Sea. Mesocosm were either manipulated to be colonized by macrophytes, or stayed phytoplankton dominated. Oxygen evolution was monitored over a period of five months in 5 min (mesocosms) to 10 min (ecosystem) intervals. Surface and depth-integrated production was calculated to analyse seasonal and areal resolved production patterns. It was found that macrophyte mesocosms were more stable, when considering only surface O_2_ production. However, calculating depth-integrated production resulted in net-heterotrophy in both shallow mesocosms approaches and the actual ecosystem. This heterotrophy is likely mediated by sediment respiration and POC accumulation in mesocosms, and a low share of productive to respiring water column in the actual ecosystem. Therefore, it seems unlikely that macrophytes will re-settle, as constant net-heterotrophy may allow for high-nutrient turnover at sediment-water interfaces and within the water column, favouring phytoplankton. Changes within the ecosystem cannot be expected without further restoration measures within and in the systems proximity.

## Introduction

### Oxygen depletion caused by eutrophication and its consequences

Transitional water bodies such as lagoons or estuaries are prone to eutrophication due to nutrients received from their catchment area. Lagoon or estuary characteristics such as spit formation can alter the water exchange rate, trapping nutrients within the water column for a long period of time [1]. One major impact of eutrophication on transitional water bodies is a change of alternative stable states from macrophyte- to phytoplankton-dominated primary production [2]. Usually, macrophyte-dominated systems produce enough oxygen to prevent suboxic conditions in the water body. In contrast, phytoplankton dominance of aquatic systems is associated with high turbidity and light limitation [3,4]. This reduced light availability decreases the euphotic depth, that means the net-autotrophic water column, which can reduce the habitable zone for submerged macrophytes, as well as light-limiting the phytoplankton itself. This decrease lowers the overall primary production per m^−2^ of the water body, even in shallow waters. Especially high phytoplankton biomasses can lower oxygen availability during night times or times with low photosynthetic active radiation (PAR), due to decreased production but still high respiration. The subsequent following suboxic conditions have major impacts on abiotic and biotic matter fluxes. For example, low oxygen concentrations lead to nutrient refluxes from sediment of phosphorus (P), as iron is reduced and insoluble FePo_4_ concentration decreases [5]. Simultaneously, zooplankton and other macrozoobenthos suffer from suboxic conditions, which lead to death and overall lowered grazing rates on phytoplankton. To complicate the issue, eutrophication-influenced sediment can show high amounts of organic carbon [6]. Such carbon is respired and consumed by bacteria and macrozoobenthos. Therefore, sediment can contribute to lags in system improvement [7], as the respiration is increased within the system. Increased nutrient recycling and supplies at reduced grazing control causes high phytoplankton biomass and can trap such eutrophic system within a phytoplankton dominated state, as phytoplankton turn-over and generation times are higher compared to submerged macrophytes.

Management of transitional waters tries to overcome these feedback mechanisms through restoration measurements such as nutrient reductions, food-web alterations, or planting submerged macrophytes [8–10]. If restoration measures lower phytoplankton primary production, the diminished macrophytes may not be able to compensate for lowered oxygen production. This lack of oxygen supply would again stabilise phytoplankton dominance as part of the above mentioned feedback mechanism. Hence, restoration of transitional water bodies might fail where phytoplankton biomass is not low enough for submerged macrophytes to come to dominance. However, in shallow margins of such transitional waters, where light is not limiting, macrophytes and phytoplankton can co-occur. The question arises why macrophytes do not gradually expand their canopy from sites with dense populations into more and deeper regions. It is possible that phytoplankton production is increased in such shallow areas as well, due to nutrient run-off [11]. Despite the possibility for macrophytes to grow, they are still hampered due to the still increased phytoplankton growth at the margins. Mechanisms may not be apparent within the entire water body, and are overlooked, due to horizontal and vertical mixing. We assume that shallow margins of eutrophic lagoons and estuaries are prone to this transient condition, as light may not be limiting, and macrophytes co-occur with phytoplankton. Experimental approaches such as mesocosms can be used to define the balance between primary production of macrophytes and phytoplankton [12,13]. This balance of production needs to be measured in transitional waters under eutrophication where organic and nutrient content in the sediment is increased. Increased knowledge of production patterns and potentials in shallow waters can thereby improve water restoration measures.

### The Darß-Zingst lagoon system – a system with failed restoration measures?

The Darß-Zingst lagoon system (DZLS), a typical model ecosystem for cold temperate lagoons of the southern Baltic Sea, lost their dense macrophytes stands 40 years ago [14,15]. Even after 30 years of reduced nutrient inputs from point sources, the macrophyte stands were not able to recover and re-colonise densely. This outcome is surprising as nutrient loads were reduced by up to 80% for phosphorus and nitrogen [16]. However, recent studies showed that the diffuse nutrient inflows across the land-water contact zone can stimulate phytoplankton growth [11,17], making shallow areas possible highly productive areas in turbid, eutrophic lagoons. If nutrients are not limiting, the other key drivers of primary production are light and temperature.

The aim of this study was therefore to investigate and model primary production of phytoplankton and submerged macrophytes in shallow water bodies. *Ex situ* mesocosms were used for primary production assays to investigate production of phytoplankton and submerged macrophytes over the duration of one growth season (May – October). Dissolved oxygen concentrations were logged continuously in six mesocosms with and without macrophytes and within the adjacent lagoon system at one point. The assay ran through one growth season to determine possible effects of temperature and light on primary production in shallow areas. We expected a seasonal difference in phytoplankton light adaptation revealed by photosynthesis-irradiance curve parameters between the mesocosms and open water of the DZLS. Further, we expected that primary production of macrophytes, once established, would increase even during phytoplankton blooms. It was hypothesized that production of submerged macrophytes would buffer short-term fluctuations of phytoplankton production caused by sudden changes in light availability or temperature and thereby prevent oxygen depletion within the water column. Difference in light adaptations and photosynthesis of phytoplankton between shallow and open waters should provide information on whether the DZLS is in a slow transition towards dominant macrophyte production or is instead blocked at the current phytoplankton community by feedback mechanisms, like photo- or nutrient acclimation. The results are discussed in the context of possible restoration measures for shallow lagoons in intermediate phases during the recovery from eutrophication.

## Material and Methods

### Description of the study site

The DZLS covers an area of 197 km^2^, and has a total volume of 397 10^6^ m^3^, and has a catchment area of 1600 km^2^ [18]. There is a strong salinity gradient from the west to east which depends upon the inflow of water from the Baltic Sea at the eastern end. The annual phytoplankton biomass (biovolume of 3.6 to 27.9 mm^3^ l^−1^) and production (109 to 611 C m^−2^ a^−1^) follows a productivity gradient from west (lowest salinity, freshwater) to east (highest salinity, brackish) pointing out a trophic status from polytrophic to slightly eutrophic [1].The experiments were conducted in the central part of the lagoon system at the Zingster Strom (ZS). The ZS is subject to changing in- and outflow events between eutrophic to polytrophic lagoon basins.

### Mesocosm detailed description

The primary production assay mesocosms were cylindric plastic containers with a height of 55cm and a diameter of 65 cm. They stood at the premises of the Biological Station Zingst. Six mesocosms were put 40cm into the ground for temperature control through cooling by the surrounding soil. Mesocosms were built up and filled with sediment and water from the ZS in March 2014. The ZS has an automatic monitoring system for oxygen, pH, temperature and conductivity. Data management and sensor maintenance were conducted by the Biological Station Zingst (University of Rostock) which is in close proximity (<50 m). Each mesocosm was filled with 0.0165 m^3^ sediment creating an approximately 5 cm thick sediment layer on the bottom. The sediment originated from a spot close to the ZS that is one of the only spots densely populated with submerged macrophytes, mostly *Chara* spp. Water from the ZS was pumped into the mesocosm until each mesocosm contained 120 l brackish water. The phytoplankton community species composition of the DZLS was stable during recent years with a permanent dominance of cyanobacteria [1,19,20]. Dominant species in these studies were by biovolume and cell count small cyanobacteria of the genus *Synechococcales*, either as solitary cells, trichomes or colonies. The plankton compositions in the mesocosms were controlled on a relative abundance several times during the experiment between May and October 2014. The water column was continuously stirred (Co. Neptun) to simulate water currents, induced water circulation thereby maintaining the non-stratified character of this lagoon system. The total capacity of the pumps was 250 l h^−1^, so that the whole water column was mixed twice an hour. Mesocosm walls and pumps were cleaned on a regular basis to remove epiphytic growth. All mesocosms were regularly checked for macrophytes. Three of them were harvested every time macrophytes were detected (phytoplankton mesocosms). The other three were left unharvested until the final sampling in October (macrophyte mesocosms). Days upon which manipulating or cleaning of the mesocosm were conducted were later removed prior the data analysis.

### Abiotic and biotic parameters

Abiotic parameters included dissolved inorganic phosphate (DIP), dissolved inorganic nitrogen (DIN, sum of nitrate, nitrite and ammonium), turbidity (absorption at 720nm), cyclic dissolved organic matter (cDOM, *gelbstoffe*, absorption at 380 nm). Biotic parameters included Chlorophyll *a* (Chl *a*, proxy for phytoplankton biomass). Chl *a* and turbidity were determined once a month (May to July) and at least biweekly from August to September. Additionally, the same set of above described abiotic and biotic parameters were sampled daily at 08:00 (CET) in the ZS as part of the daily monitoring at the BSZ. A final sampling in the mesocosms was conducted in October, including sampling of submerged macrophyte biomass. The sampled biomass was separated into species, and wet weighed. Macrophyte biomass was dried at 90 °C for 24 h in an oven and dry weighed again afterwards. The macrophyte dry mass was further processed by grinding, weighing in tin capsules, and adding one drop of HCl to volatilize inorganic carbon. Finally, macrophyte samples were measured in an elemental analyser (Vario-EL III) for organic carbon and nitrogen. Additionally, at two occasions samples were taken to determine seston and particulate organic carbon and nitrogen in the water column. Particulate organic carbon (POC) and nitrogen (PON) of seston was determined using by harvesting particles from the water onto pre-combusted glass fibre filters (GF/F, Whatman). Filters were then treated in HCl vapor over night to volatilize inorganic carbon. Afterward particle filtration the filters were measured in an elemental analyser (Vario-EL III) for carbon and nitrogen. The elemental analyser was calibrated with acetanilide. All values from abiotic parameters are presented as monthly medians.

### Mesocosm and ecosystem community production

Monitoring of oxygen evolution begun in the mesocosms after the first sprouted macrophytes were detected in May. Dissolved oxygen concentration and temperature were recorded every 5 min from mid-May until mid of October 2014 using optode loggers (Hq40, LDO, Hach-Lange) 5cm under the surface in each mesocosm. The dissolved oxygen concentrations in the ZS were recorded every 10 min continuously at a water depth of 50 cm (total water depth 5 m) for the entire year of 2014. Oxygen evolution in the ZS was measured at the same spot the monitoring program of the abiotic and biotic parameters. Percent oxygen saturation measured was converted to mmol O_2_ l^−1^ using solubility values for the brackish water of the mesocosm and the ZS [21] considering temperature and salinity.

The diffusive fluxes into the mesocosms and the ZS were compensated using the formula of D’Avanzo et al. [22] due to over- and undersaturation of O_2_ in relation to temperature and salinity. Additionally, the gas exchange coefficient for the mesocosms was empirically determined in one additional mesocosm, which was driven to 20% undersaturation through rapid temperature changes compared to the solution equilibrium. The kinetics of oxygen concentration were measured (optode logger, Hq40, LDO, Hach-Lange) within the same salinity and temperature range as the experimental mesocosms. The gas exchange coefficient was drawn from the slope of the established saturation curve.

Hourly rates of O_2_ evolution were calculated by regressing dissolved oxygen concentration against time and correcting for diffusion. Hourly rates over each day were summed up and regarded as daytime production expressed in mmol O_2_ l^−1^ d^−1^. Daytime production is attributed to the net community production in excess of community respiration during hours of sunlight. Daily negative hourly rates represent the night time respiration recorded in mmol O_2_ l^−1^ d^−1^. The rates and production calculated for the individual mesocosms were averaged for the two treatments with and without macrophytes.

### Photosynthetic-irradiance curves

The continuously recorded changes in O_2_ concentration were scaled to phytoplankton Chl *a* concentration per litre and reported as hourly chlorophyll-specific oxygen production rates (μmol O_2_ mg Chl *a*^−1^ h^−1^). For each day, hourly production rates were regressed against irradiance to fit photosynthesis-irradiance curves. The solar irradiance data were provided from the Biological Station Zingst and the Deutscher Wetterdienst (German Meterological Service) from their Warnemünde (St.-Nr. 04271) and Arkona (St.-Nr. 00183) stations (Figure 1). When the Zingst sensor failed for several weeks, the German Meterological Service-sensor data was used instead. Underwater irradiance (*PAR*_*0*_) was calculated from solar irradiance, sun elevation and water attenuation [23]. The water attenuation coefficient (Kd) was calculated using the formula of Xu et al. (2005):

**Figure 1:**
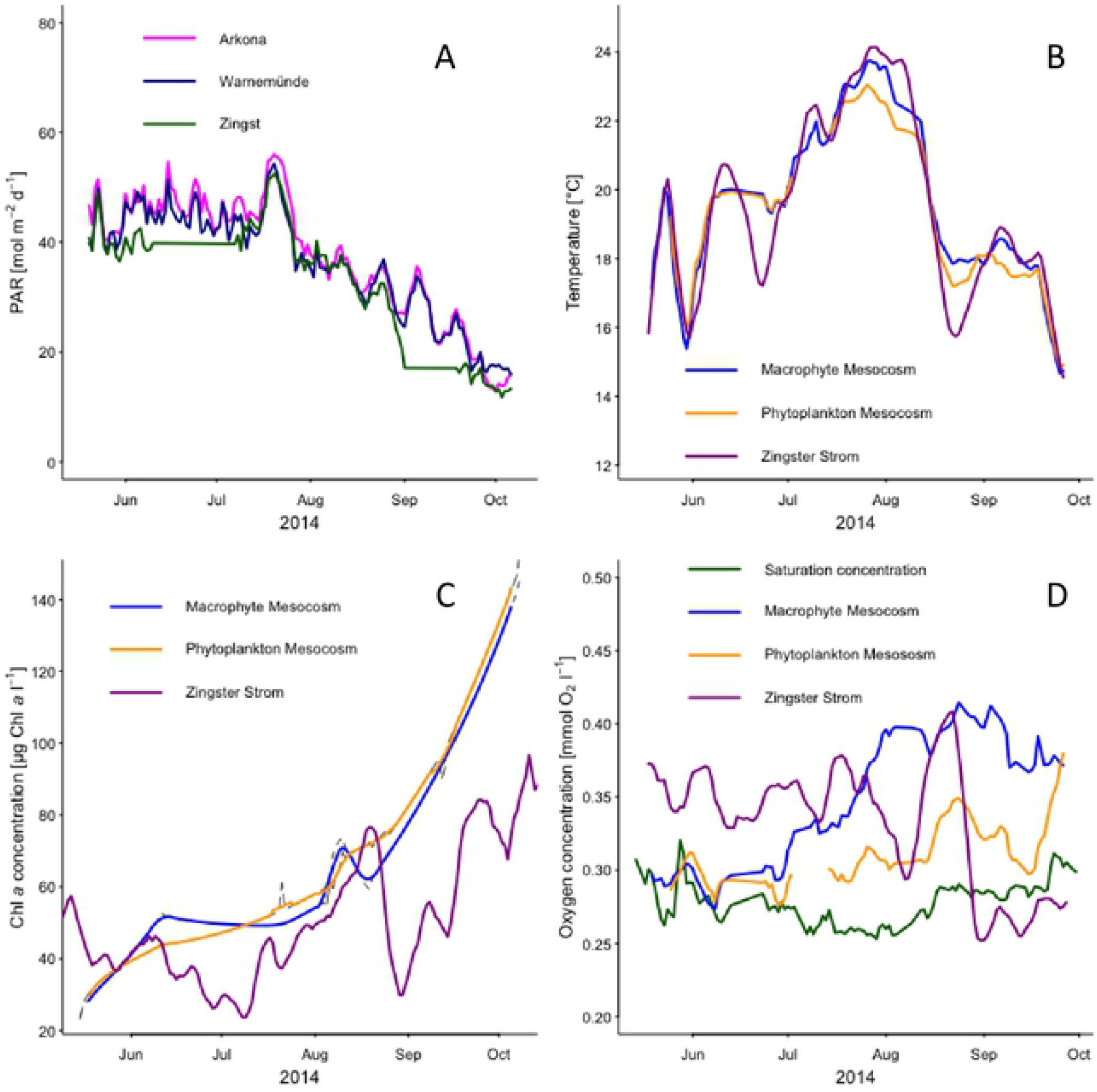
Time series of photosynthetic active radiation (PAR), temperature, Chlorophyll *a* (Chl *a*) and oxygen concentration. PAR and Chl *a* concentration, temperature and oxygen concentration are shown as daily average. Solid lines represent weekly running means fitted to oxygen concentration in mesocosm with (blue line) and without (orange line) macrophytes and the Zingster Strom (magenta line). The green line represents the oxygen saturation concentration.

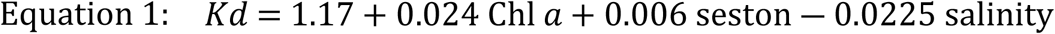

where Chl *a* is the concentration of Chlorophyll *a*, seston the concentration of suspended particular material and salinity. The irradiance at logger depths (PARz) was calculated from the irradiance data by applying the Lambert-Beer law:

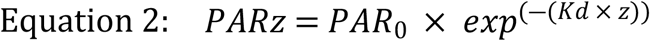

where *PAR*_*0*_ is the surface radiation, *Kd* is the attenuation coefficient based on Equation 1 and z the water depth at logger position in the water column. Depth *z* was corrected for the daily-recorded changes in water level of the ZS, but stayed the same in mesocosms.

The equation to regress hourly oxygen production *P* against *PARz* was chosen from [23]:

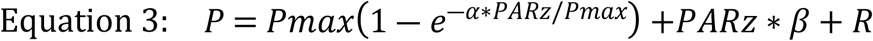

where *PARz* is the light intensity at logger depth, *Pmax* the maximum oxygen evolution rate, *α* the initial slope of the production curve, *Β*the slope at high irradiance due to photoinhibition, and *R* the hourly respiration rate. Mesocosms and ZS community respiration rates were derived from average hourly night time respiration rates and reported as hourly chlorophyll-specific oxygen respiration rates (μmol O_2_ mg Chl *a*^−1^ h^−1^). Separate curves were fitted for each day of measurement in each mesocosm to estimate daily photosynthetic parameters (*P*_*max*_, *α, β*) and respiration rates.

The initial photosynthetic parameters (*P*_*max*_, *α, β*) were taken from Schumann et al. [24]. The margins of this parameter were corrected for water temperature difference between the original study and the mesocosm [25]:

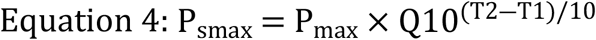

*P*_*smax*_ is the seasonal maximum production at T2 observed water temperature in the mesocosm, *P*_*max*_ the maximum production at temperature T1 and the Q10 factor. The nonlinear least square estimates of the photosynthetic parameters were determined using the *multiStart*-function in the BB package [26] in the R software environment (R core team 2014). The photosynthesis light saturation point (*I*_*k*_) was calculated as the quotient between *Pmax* / *α* [27]. The compensation light intensity (*Ic*) was derived from equation 3 to calculate the compensation depth using equation 2, where respiration demands equal photosynthesis oxygen production. Averages of all calculated daily photosynthetic parameters were grouped for the ZS, phytoplankton and macrophytes mesocosms and reported for the temperature intervals below 17°C, 17-20°C, 20-23°C and above 23°C (Table 1).

**Table 1.** – Photosynthetic production vs. irradiance curve parameters (± standard error) based on the Walsby [21] equation fitted to the data and averaged for four different temperature ranges. *∝* initial slope of the light saturation curve μmol O_2_ mg Chl *a*^-1^ h^-1^ (μmol photons m-^2^ s^-1^)^-1^; *P*_*max*_the maximum production rate μmol O_2_ mg Chl *a*^-1^ h^-1^; *I*_*c*_: light compensation point (μmol photons m-^2^ s^-1^); *I*_*k*_: point where the linear initial slope intersects with the light intensity of *P*_*m*_ [27] (μmol photons m^-2^ s^-1^), to denote the onset of light saturation.

| Photosynthetic<br>parameters | Temperature |  |  |  |
| --- | --- | --- | --- | --- |
|  | <17°C | >17°C <20°C | >20°C <23°C | >23°C |
| Macrophyte mesocosm |  |  |  |  |
| $\alpha$ | 1.9±0.2 | 1.8±0.2 | 1.7±0.1 | 1.6±0.5 |
| $P_{max}$ | 319.7±10.1 | 362.3±7.5 | 453.1±7 | 472.8±10.7 |
| $I_c$ | 84.7 | 96.8 | 154.4 | 130.9 |
| $I_k$ | 285.0 | 300.4 | 353.3 | 274.8 |
| Phytoplankton mesocosm |  |  |  |  |
| $\alpha$ | 2±0.3 | 2.1±0.1 | 2±0.2 | 2.1±0.3 |
| $P_{max}$ | 304.1±11.8 | 383.9±5.1 | 471.8±7.7 | 530±8.1 |
| $I_c$ | 78.1 | 91.6 | 153.8 | 151.9 |
| $I_k$ | 310.4 | 236.5 | 313.4 | 284.7 |
| Zingster Strom |  |  |  |  |
| $\alpha$ | 3±0.1 | 3.2±0.1 | 3.5±0.1 | 3.2±0.1 |
| $P_{max}$ | 285.1±2.6 | 333.0±3.3 | 382.9±3.0 | 422.8±2.1 |
| $I_c$ | 31.2 | 40.2 | 53.6 | 56.2 |
| $I_k$ | 95.0 | 106.1 | 110.6 | 131.1 |

### Depth integrated vs. surface community production

We used calculated volumetric daytime production and night time respiration to calculate the surface daily net community production. It is the sum of the latter two rates representing the amount of O_2_ produced at the surface during the hours of sunlight, which was not consumed during the following night. Surface daytime gross community production of the mesocosm and the ZS was the time integrated sum of net daytime production and a daytime respiration demand, the latter being calculated from the average night time mesocosm respiration rate multiplied by the hours of daylight. The surface production only represents productivity patterns around the optodes.

In a second approach hourly chlorophyll-specific oxygen production rates (μmol O_2_ mg Chl *a*^-1^ h^-1^) were calculated separately at 0.1m depth steps in each mesocosm and the ZS using the underwater irradiance (PARz) and estimated daily photosynthetic parameters (see above). Hourly community production rates (μmol O_2_ l^-1^ h^-1^) were the product of the hourly chlorophyll-specific oxygen production rates and chlorophyll concentration at each depth step and time interval. The depth integral of the hourly community production (μmol O_2_ dm^-2^ h^-1^) at a given time in each mesocosm was obtained by the trapezoidal rule [23]. Daily integrals of the depth integrals of the hourly community production (μmol O_2_ dm^-2^ d^-1^) were calculated in the same way and represent the daily net community production. Daily net community production was the amount of O_2_ produced in each mesocosm and the ZS during the hours of sunlight, which was not consumed during the following night. Daytime gross community production of the mesocosm and the ZS was the time integrated sum of net daytime production and a daytime respiration demand, the latter being calculated from the average night time mesocosm respiration rate multiplied by the hours of daylight and number of integrated depth steps. Daily net and gross community production were scaled to one square meter and reported as mmol O_2_ m^-2^ d^-1^ to allow the comparison between the mesocosms, the ZS and literature values.

## Results

### Environment parameters in mesocosm and the Zingster Strom

Average daily photosynthetic active radiation (PAR) was stable from May to mid July 2014 (Figure 1). There was a long phase of high PAR corresponding with a water temperature increase in mesocosms and ZS (Figure 1B), which was also the peak during this year. The temperature of the mesocosm followed the temperature of the ZS within ±1 °C (overall median of daily temperature difference, n=493) (Figure 1).

In all mesocosms, pH stayed constant at 8.9 ± 0.5. The concentrations of DIN and DIP were always very low in all mesocosms. DIP never exceeded 0.3 μmol l^-1^ (August), which was the only occasion where DIP was above the quantification limit of 0.05 μmol l^-1^. NH_4_ contributed the largest fraction of DIN with up to 6.2 μmol l^-1^ (with a median at 2 μmol l^-1^), and NO_3_ and NO_2_ were below 0.05 μmol l^-1^ 90% of the time (n= 4 – 6 per mesocosm). Monthly medians of the ZS (daily measurements) were between 0.7 – 3.7 μmol DIN l^-1^ and 0.0 – 0.15 μmol DIP l^-1^ with highest concentrations in June, and lowest in August for both nutrient fractions (see Supplement Table 1).

The salinity was stable from May until July (7 PSU) in all mesocosms and decreased in August (6 PSU) and did not change until October. In ZS, lowest salinity was measured in April (6 PSU) and highest in September (7 PSU). The Chl *a* concentration was 85 μg Chl *a* l^-1^ at the start date (March 2014, data point not shown) in all mesocosms and the ZS. It decreased until May in the mesocosms to a median of 23 μg l^-1^. Chl *a* concentration in the ZS decreased in the same time to 42 μg Chl *a* l^-1^, indicating the same trends within mesocosms and the actual ecosystem. Chl *a* concentrations peaked with 136 μg l^-1^ in the mesocosms and 95 μg l^-1^ in the ZS in October. The seston concentration almost doubled within the mesocosms from March (45 mg l^-1^) to August (70 mg l^-1^), and further increased until October (84 mg l^-1^). In contrast the ZS showed monthly fluctuating seston medians ranging from 65 mg l^-1^ (May) to 23 (October). Accordingly, the attenuation coefficient changed from 1.9 to 3.4 (macrophyte mesocosms, range 0.9 – 4.1) and 4.5 (phytoplankton mesocosms, range 1 – 7.3) from filling up in March to October. The attenuation coefficient in the ZS ranged from to 1 to 4.8 with lowest values found in July and two peaks in August and October.

The O_2_ concentration exceeded the O_2_ saturation concentrations in both mesocosm treatments from June until October (Figure 1) with only two short periods of undersaturation indicating intervals of system net-heterotrophy. From June onwards, the O_2_ concentration was higher in mesocosms with macrophytes. The ZS showed O_2_ concentrations above saturation level until end of August, when a sudden decline appeared. The O_2_ concentration stayed below saturation concentration until end of observations in October.

### Elemental composition and development of primary producers

Chl *a* concentrations in macrophyte mesocosms were not significantly lower, compared to phytoplankton mesocosms. Relative community abundance in the mesocosms showed the same species distribution as the ZS over the time (personal observation). Particulate-bound elements increased over time and peaked in October with up to 58 mg l^-1^ (POC) and 5.7 mg l^-1^ (PON) showing an accumulation of elements within mesocosm waters. Simultaneously, the ratio of POC: seston (average ratio 0.25 mg POC: 1 mg seston, n = 8) and POC: Chl *a* (average ratio 234 μg POC: 1 μg Chl, n = 8) remained constant, indicating a stable increase of phytoplankton as part of seston. Unfortunately, there are no comparable POC and PON values from ZS for 2014.

Sprouting of macrophytes started in all mesocosms in May until July and stopped from August until September. One of the repeatedly harvested phytoplankton mesocosm showed macrophyte biomass again in October, indicating a second sprouting period in September/October. Macrophyte biomass and POC in macrophyte mesocosms was three times as high (150 mg DM m^-2^, 43.4 mg POC m^-2^) as the sum of all macrophyte harvests in phytoplankton mesocosms (56 mg DM m^-2^, 14.6 mg POC m^-2^, see Supplement Figure 2). Among the macrophyte species were *Najas marina, Chara baltica*, and *C. hispida*, with charophytes dominating the biomass representing 90% of the dry mass.

**Figure 2.**
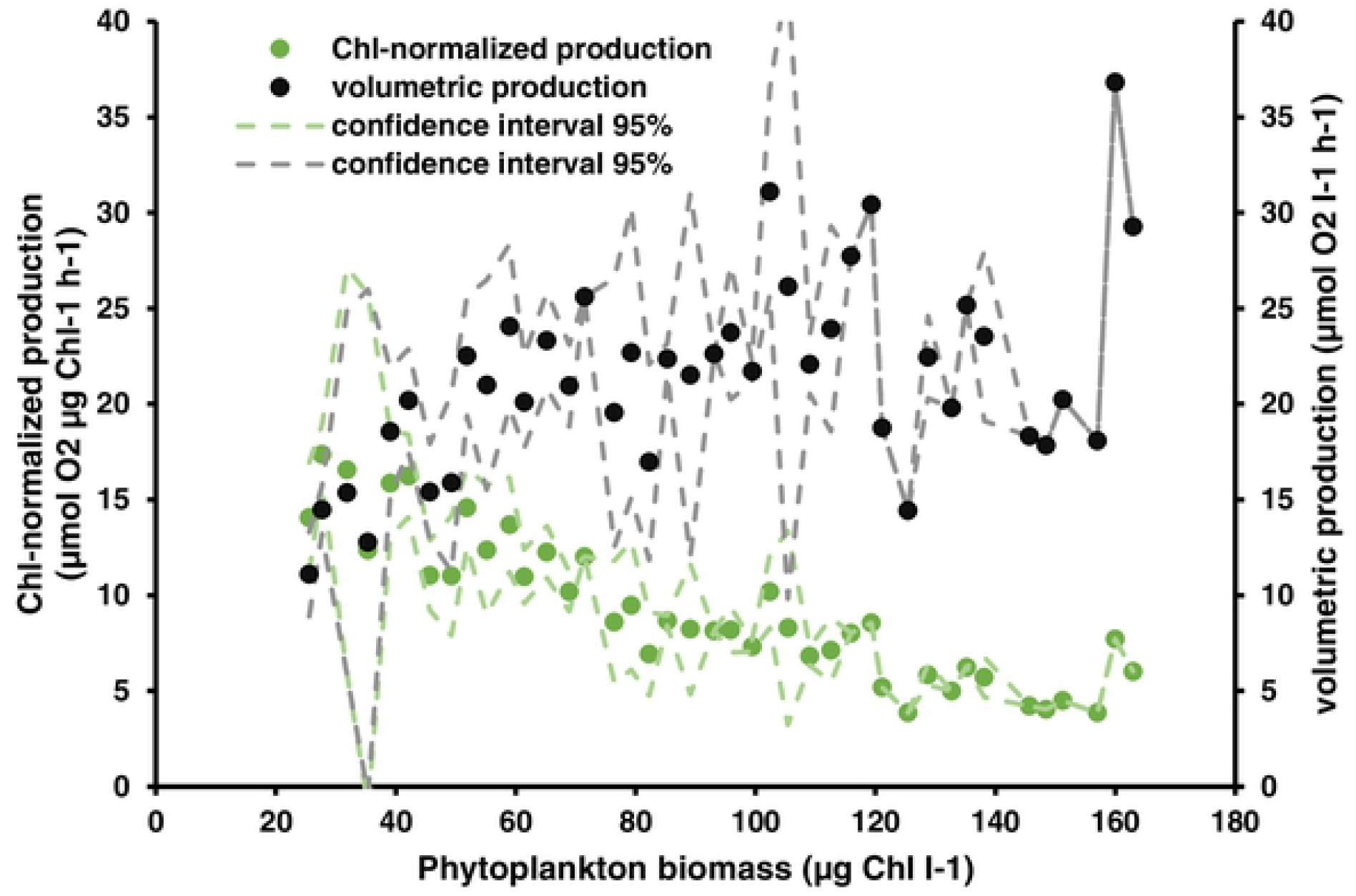
Chl-normalized (μmol O_2_ μg^-1^ Chl *a* h^-1^) based on phytoplankton biomass (Chlorophyll *a* μg l^-1^) and volumetric (μmol O_2_ l^-1^ h^-1^) hourly oxygen production in phytoplankton mesocosms. Production values represent averaged hourly values calculated from daily production values divided by hours of daylight. Confidence intervals are based on standard deviations of pooled, averaged Chl-concentrations (interval steps at 3 μg L^-1^).

### Daytime production response to ambient irradiance and temperature

Macrophyte mesocosms showed consistently lowest *α* values, compared to phytoplankton mesocosms and the ZS (Table 1). The maximum production rates (*P*_*max*_) of the fitted P/I curves peaked at water temperature above 23°C in both mesocosm treatments, as well as in the ZS. Both mesocosm treatments showed higher *P*_*max*_ values at all temperatures than the ZS. On days falling below this temperature range, *P*_*max*_ and the compensation light intensity (*I*_*c*_) were at lower levels in the mesocosms and the ZS. Both mesocosm treatments showed equally high *I*_*c*_, which were up to three times higher compared to the ZS. Light intensities to reach light saturation was highest in macrophyte mesocosms, being up to three times higher compared to the ZS.

### Surface net and gross community production

Averaged volumetric hourly O_2_ production (μmol O_2_ L^-1^ h^-1^) tended to increase with increasing phytoplankton biomass up to 60 μg Chl *a* L^-1^ in phytoplankton mesocosms (Figure 2). However, O_2_ production remained on a plateau of around 20 μmol O_2_ L^-1^ h^-1^, and possibly decreased again at very high phytoplankton biomasses (>100 μg Chl *a* L^-1^). Normalizing this production to the respective Chl *a* values showed a constant decrease of O_2_-production per Chl *a* with overall increasing biomass. This pattern indicates increasing light limitation, even though volumetric production remained high.

Daily surface net community production ranged between −0.04 and 0.12 mmol O_2_ l^-1^ d^-1^ in macrophyte mesocosms and between −0.09 and 0.18 mmol O_2_ l^-1^ d^-1^ in phytoplankton mesocosms. Surface net community production peaked in both treatments at the end of May and fell below zero at the beginning of June and subsequently increased to maximum levels in July and August (Figure 3 A). Surface net community production in phytoplankton mesocosms exceeded surface net community production of macrophytes mesocosms in beginning of July and August, and at the end of September. Macrophyte mesocosms showed only one sharp net production drop in September. Phytoplankton mesocosms showed an overall greater variability than macrophyte mesocosms with drops in surface net production several times within the observed time. In contrast, surface gross community production reached its maxima in mid-July with macrophyte mesocosms always having a higher surface gross community production compared to phytoplankton mesocosms (Figure 3 B).

**Figure 3.**
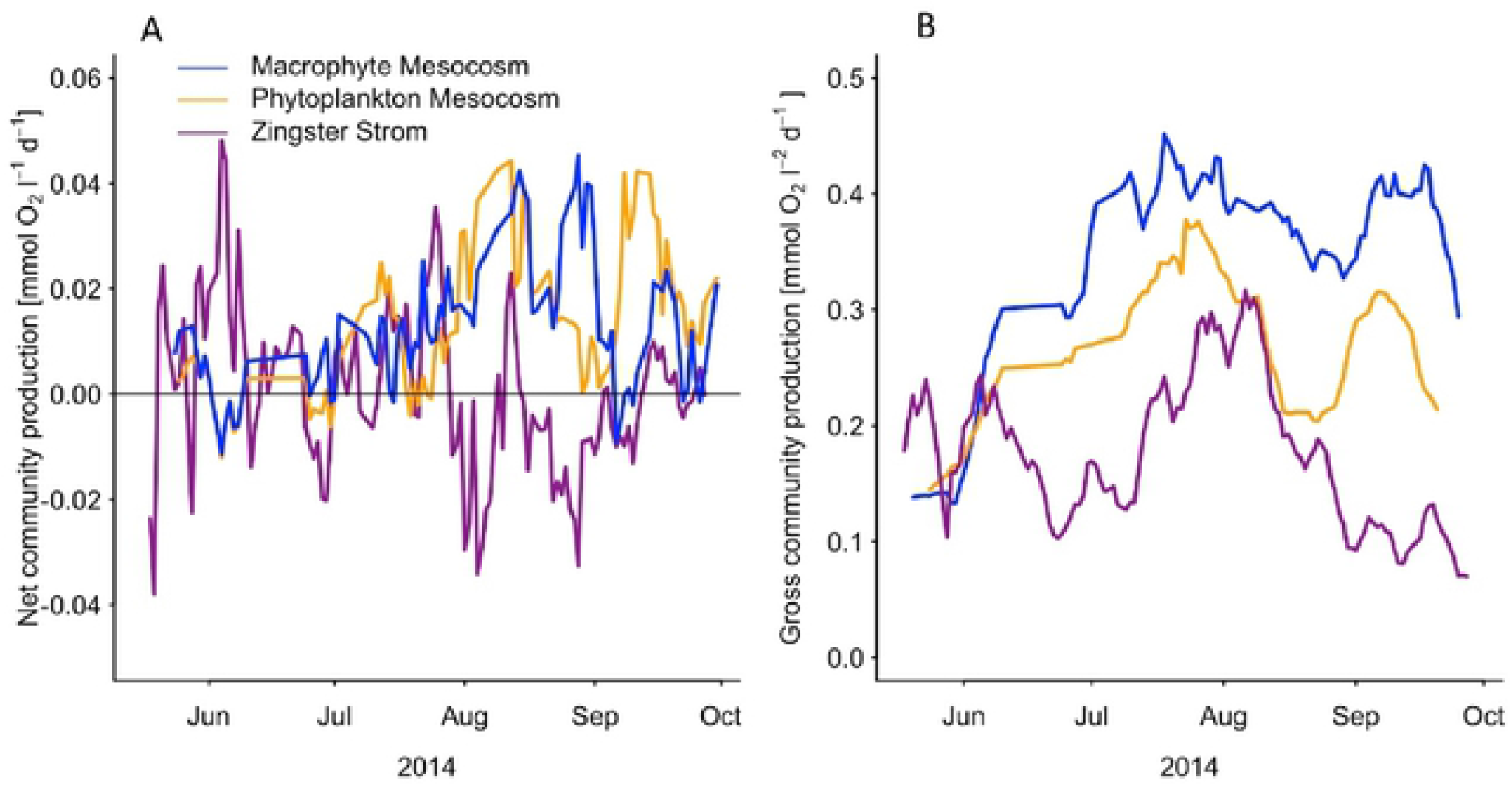
A – Surface net community production (mmol O_2_ l^-1^ d^-1^), the sum of daytime production and night time respiration, in macrophyte mesocosms (blue), phytoplankton mesocosms (orange), and the Zingster Strom (magenta). Dates of cleaning and sampling including the following day were removed prior the analysis. Weekly running mean of daily values are shown. B – Surface daytime gross primary production, in macrophyte mesocosms (blue), phytoplankton mesocosms (orange) and the Zingster Strom (magenta), calculated as the integrated sum of net daytime production and a daytime respiration demand. Daytime respiration was calculated from the average night time respiration rate of the respective system and multiplied by the hours of daylight. Values represent only the production around the sensor at the respective depths.

The presence of macrophytes in the mesocosm had a positive effect on the gross community production of the mesocosms. Beside a short period at the beginning of the experiment, the surface gross community production in macrophytes mesocosm was on average higher than in phytoplankton mesocosms. Phytoplankton mesocosms experienced a larger drop in surface gross production during August, compared to macrophyte mesocosms. The seasonal trend was similar as for daily surface net community production, starting with low values in June, increasing to the maximum in July and August and declining towards the end of the experiment. However, surface gross community production decline was more pronounced in phytoplankton mesocosms.

The daily surface net community production of the ZS ranged between −0.3 and 0.4 mmol O_2_ l^-1^ d^-1^ (Figure 3A). Although daily surface net community production showed strong day to day variation, the weekly running mean stayed mainly positive from the beginning of June until the end of August. The daily surface gross community production ranged between 0.06 and 0.3 mmol O_2_ l^-1^ d^-1^ and showed highest values in August. Both, daily surface gross community production and daily surface net community production, declined strongly at the end of August and daily gross community production stayed low until the end of the observation period.

### Depth integrated community production

The areal daily net community production ranged on average between −837 and 613 mmol O_2_ m^-2^ d^-1^ in phytoplankton mesocosm and between −469 and 455 in macrophytes mesocosm. Weekly areal daily net community production average was negative over most of the time and only indicated autotrophy end of May and beginning of July in both treatments (Figure 4A). Macrophyte mesocosms showed less variability in daily net community production. Both treatments daily net community production decreased from July until mid of September. Only macrophyte mesocosms became net autotroph in beginning of October. The areal daytime gross community production showed ranges between −129 and 759 mmol O_2_ m^-2^ d^−1^ in phytoplankton mesocosms and −56 and 964 mmol O_2_ m^−2^ d^−1^ in macrophytes mesocosm. From June onwards macrophytes mesocosm had a lower daytime gross community production. Both treatments reached their highest daytime gross community production mid of July, which decreased afterwards until September before increasing again until end of September.

**Figure 4.**
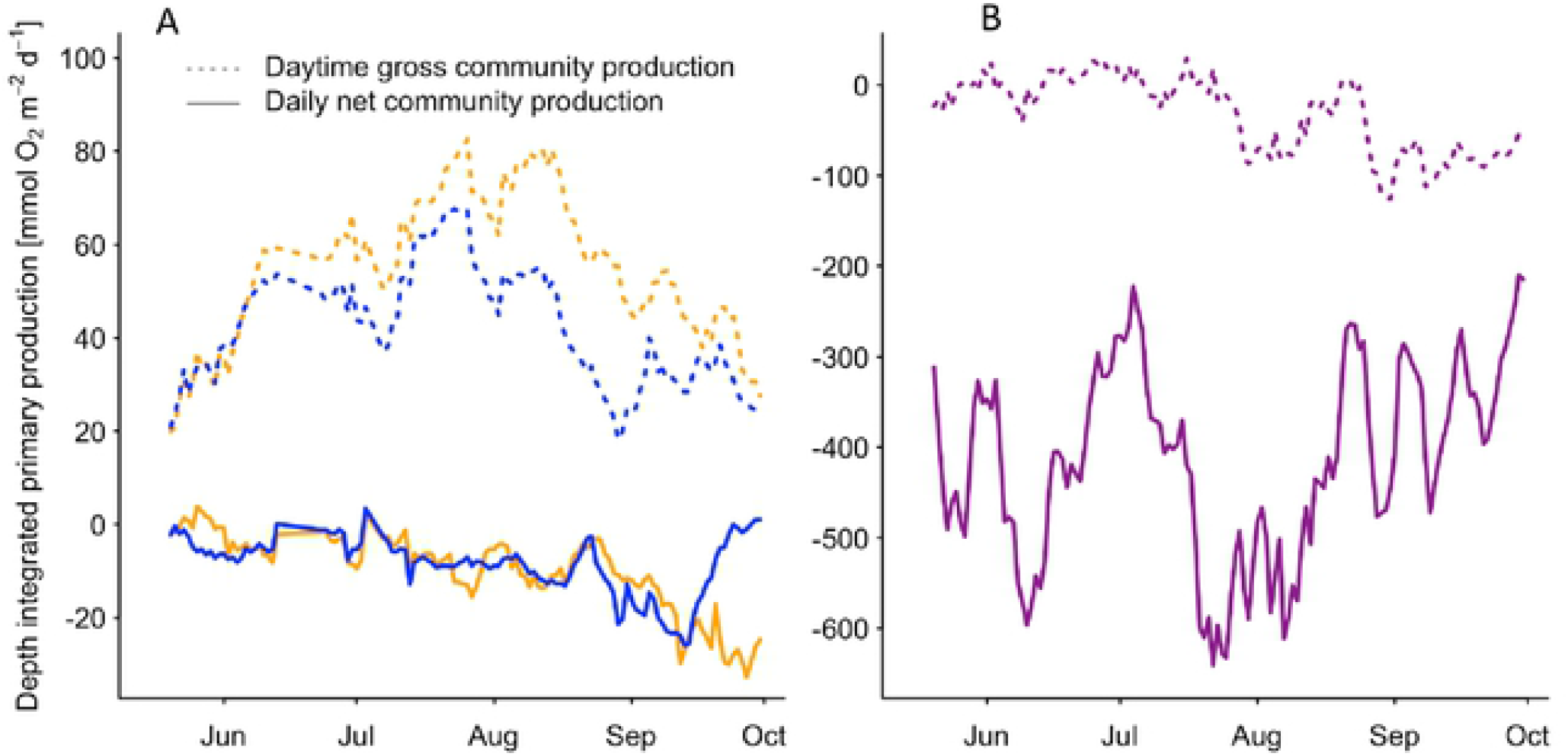
A – Depth-integrated daily net community production (mmol O_2_ m^-2^ d-1), the sum of daytime production and night time respiration, and daytime gross community production (mmol O_2_ m^-2^ d^-1^), calculated as the integrated sum of net daytime production and a daytime respiration demand, in macrophyte mesocosms (blue), and phytoplankton mesocosms (orange). Daytime respiration was calculated from the average night time respiration rate of the respective system and multiplied by the hours of daylight. Dates of cleaning and sampling including the following day were removed prior the analysis. Weekly running mean of daily values are shown. B Depth-integrated daily net community production and daytime gross community production in the Zingster Strom (magenta) calculated as stated above. Please note the different scaling.

In the ZS the areal daily net community production was one magnitude lower per square meter in comparison with both mesocosm treatments and ranged on average between - 11167 and −415 mmol O_2_ m^-2^ d^-1^. Zingster Strom daily net community production had two minima in beginning of June and end of July and showed highest values in mid July and beginning of September. Daily net community production showed highest variability in September. The areal daytime gross community production had a range between −2771 and 1847 mmol O_2_ m^-2^ d^-1^ and showed highest values in June and July. Its weekly average decreased until October.

## Discussion

### Methodological annotations

The mesocosms ran for almost six consecutive months exposed to the same environmental conditions as the adjacent lagoon system. However, the small volume, and enclosed water body without water exchange led to some alterations in hydro-physical and -chemical composition. For example, water temperature offset was around ±1 K on median compared to the actual system. Furthermore, dial deviation reached up to 4 °K offset from the ZS and depended strongly on the daily radiation. The median temperature difference was comparable to other land-based mesocosm approaches, like actively cooled benthocosms of Sylt [29], and Kiel [30] with ±1.5 K on median. Nonetheless, this temperature deviation had likely an effect on primary production (see below). Dissolved nutrients were always very low (< 0.3 μmol l^-1^ for DIP, < 6.2 μmol l^-1^ for DIN) within the mesocosms as well as the lagoon (see supplement table 1). The low availability of dissolved inorganic nutrients is a special feature of the DZLS, probably caused by its cyanobacterial dominance [20], which were described to rapidly take up occurring nutrient pulses [31].

Water circulation by water pumps simulated the permanently mixed shallow areas of the lagoon system. The permanent mixing prevented a possible light limitation for phytoplankton even in the late phase of the experiment with highest primary producer biomass and turbidity. The mixing increased possibly primary production by evenly distribution and timing of phytoplankton within the water column [32] and can explain the deviation in oxygen concentration, but not in oxygen production, between lagoon system and mesocosms (see Figure 1, 6 and discussion below). The long run-time of this artificial situation is justified by the results, as at least 30 days are necessary to characterize variations in summer production [33]. Furthermore, sprouting and growing of macrophytes took at least two months within the mesocosms. Contrary to the actual system, exclusion of large grazers and macrozoobenthos changed the trophic food web, but allowed the determination of the planktonic community production. Both, Chl *a* and seston concentration, indicated a strong accumulation of organic material and phytoplankton in the mesocosms. Interestingly, Chl *a* concentrations increased steadily from 20 to 140 μg l^-1^ pointing to a low grazing control of phytoplankton by zooplankton. Typical daily grazing rates on phytoplankton were described to range between 0.6 to 6% of primary production going into higher trophic levels [34,35]. Nonetheless, the higher Chl *a* concentrations are sometimes found in the inner parts of the lagoon system within the last 15 years [36], indicating a shift from a meso-/eutrophic state to eu-/polytrophic state over the course of the experiment. However, Chl *a* was comparable to the actual system, but seston increased due to the shallow, enclosed water body within the mesocosms.

### Effects of temperature, nutrients, and light on primary production

Determination of net-autotrophic or heterotrophic conditions depend on defined assumptions regarding light availability and plasticity of conversion factors (i.e. O_2_ to C). Additionally, C: Chl *a* ratios of phytoplankton depend strongly on light intensity and temperature during growth and are not linear [37]. Conversions from Chl *a* to C and *vice versa* are therefore impossible with the methodology used in this study. We therefore focused on differences in O_2_ production capacities.

Temperature had a major effect on primary production, as physiological rates increase at higher temperatures [38]. The pre-dominant pico-cyanobacteria of this system were stimulated by higher temperature (see Figure 1 & Table 1, [13]). The same finding was described in Blanes Bay, where picophytoplankton showed a Q_10_ of 1.85 [39]. Similarly, we found an almost 100% decrease of primary production in phytoplankton mesocosms, when temperature dropped from 23 to 17 °C within one week in August. Macrophyte mesocosms were less affected, even though two out of three mesocosms showed lowered Chl *a* concentrations. Macrophytes likely buffered the production gap. The one macrophyte mesocosm with still increasing phytoplankton biomass had only half the macrophyte biomass than the other two. Similar depressions on phytoplankton biomass were already described for *Chara* stands in shallow lakes [40]. Interestingly, phytoplankton mesocosms were able to restore their former daytime production after that drop, even though temperature remained at 17 °C. This stabilization of production was probably related to the still increasing Chl *a* concentration within the water column (Figure 1). The increasing phytoplankton biomass produced the same total amount of oxygen, but at lower rates per Chl *a* (see Figure 2). The lower normalized rates can be accounted to either lowered temperature, or self-shading of phytoplankton. An apparent phytoplankton species composition shift was not detected, as cyanobacteria still dominated, but green algae, or diatoms were not abundant (personal observation). The same temperature-decrease most likely caused a negative daytime community production in the ZS, as Chl *a* concentrations dropped by more than 60%. Subsequently, the oxygen concentration in the ZS was even below saturation concentration (see Figure 1).

The increased primary production in mesocosms likely increased bacterial respiration [12] and nutrient recycling by bacterioplankton [41]. However, no conclusion can be drawn, as counted plankton samples are missing. Nonetheless, elevated temperature increased respirational rates of the community, that means from all plankton components, as well as from macrophytes and the sediment. Macrophyte mesocosms stayed net-autotroph during prolonged low PAR intervals in mid of July. In contrast phytoplankton mesocosms became net-heterotroph over the same time. Interestingly, macrophyte mesocosms were able to acclimatise faster to changing temperature levels, compared to phytoplankton mesocosms. This finding points at overall lower buffer capacities of the phytoplankton dominated mesocosms, as turn-over is high at simultaneously high temperature [39]. Net-primary production of macroalgae was described as less affected by temperature and light changes [42].

Light availability seemed less important, as high PAR fluxes increased net daytime production, but lowered PAR did not result in lower net daytime production in mesocosms at simultaneously high temperatures. Furthermore, PAR was only decreasing gradually over the course of the season and was therefore not abruptly changing like temperature. This result points at a strong photo-acclimatisation process for both, phytoplankton and macrophyte communities. Phytoplankton of the DZLS is described to be low-light adapted [43], as underwater light availability (1% depth) is sometimes less than 1 m [44]. *P/I*-curves based on the seasonal margins of measured dominant DZLS phytoplankton photosynthetic parameters [24] were in agreement with our data. Furthermore, the dominant picocyanobacteria in this system and our study are closely related to *Synechocystis* PCC6803 [45], which showed in pure cultures three- to six-times higher *α* and *P*_*max*_ at two- to 12-times lower Chl *a* concentrations compared to this study [46]. However, our study analysed a community net-production with simultaneously high respiration rates from other ecosystem compartments (sediment) as well, as shown with our 3- to 5 times higher daytime gross community production compared to daily net community production (see Figure 3 & 4). The results from laboratory studies and field experiment seem to be in the same order of magnitude. Interestingly, *Synechocystis* PCC6803 showed highest accumulated O_2_ production during short (15 min) light-dark changes with O_2_ production decreasing over longer light-dark intervals, pointing at an adaption to turbid, well-mixed water bodies [46]. The water column in our mesocosms was completely stirred in approximately 30 mins, whereas the circulation in deeper parts of the ZS (>1 m) may take longer. These results can explain the higher oxygen concentrations within the mesocosm, compared to the ZS (Figure 1). Average hourly O_2_ production rates found in phytoplankton mesocosms are in agreement with, for example turbid, macrophyte-free lakes, where similar rates were described [47]. The comparison of areal O_2_ production rates depend on the respective water depths and analyses frequency. For example, bentho-pelagic mesocosms containing macrophytes (*Ruppia* sp.) and pelagic mesocosms containing only phytoplankton showed negative areal daily net community productions in July, September and November in St. Andre lagoon [48], which is comparable to our findings. Simultaneously, oxygen concentrations in St. Andre lagoon were also most of the time above saturation concentrations, when considering only the volumetric production at the surface. The surprising findings of overall constant net-heterotrophy within the ecosystem and the shallow mesocosms can be explained by high amounts of waterborne POC (up to 16 mg L^-1^) and DOC (up to 12 mg L^-1^), stimulating bacterial turnover (up to 18 μg C L^-1^ h^-1^, [49]) and probably respiration. Furthermore, sediments in the ZS and elsewhere in the DZLS can show combustible organic contents of up to 20% [50], indicating another oxygen sink in the system. A strong light gradient may result in net-heterotrophy already in shallow areas. The depth-integration is therefore an important aspect when comparing phytoplankton to macrophyte systems, as production patterns can be different.

Collected macrophytes in mesocosm were probably low-light adapted, too. Macrophyte species found in the mesocosms (*Najas marina, Chara baltica, C. hispida*) were described to be part of low-light adapted populations [51]. Those species can sometimes form dense patches within the adjacent lagoon system, but show a high interannual variability [14,15]. Reported *I*_*K*_ for these populations were around 30 – 60 μmol photons m^-2^ s^-1^ and simultaneously, photoinhibition and *P*_*max*_ was described to be variable, depending on antecedent light conditions [51].

However, phytoplankton biomass increased even at lowered light and cooler conditions, as Chl *a* peaked in September and October. These results point to an accumulation process, most likely of nutrients, which increased phytoplankton biomass even at lowered light and temperature. Even though dissolved nutrient concentrations were low, total nutrients likely increased over time caused by wet and dry deposition and possibly sedimental release. Total phosphorus increase through wet and dry deposition can be accounted to roughly 2 μmol l^-1^ during the experimental time [52]. This nutrient deposition through rain and dust does not include possible impacts of terrestrial invertebrates, which is likely in an open mesocosm approach. Measured values for total nutrients in the mesocosms are missing and cannot be derived from Chl: Phosphorus ratios as these ratios are skewed within inner coastal water bodies of the southern Baltic Sea [20]. However, the concentrations for particulate organic carbon and nitrogen increased, but biomass proportions of particulate matter stayed constant, based on POC: seston (see supplement table 2). It is likely that the nutrient fluxes over time were all incorporated into biomass, as dissolved nutrient levels were low (see supplement table 1). These results indicate the accumulation process, but also the steady increase of phytoplankton within the mesocosms over time. Other mesocosm studies showed that primary production increased with increased levels of nutrient addition [53], but that very high nutrient additions lower primary production. The same pattern was probably detectable in this study as well, as Chl-normalized production tended to decrease at high phytoplankton concentrations (see Figure 2 and Figure S1). The dominating small picophytoplankton within the mesocosms and the lagoon system [1,54] are probably able to take up small nutrient concentrations very efficiently [55] and have an overall low half-saturation constant for N and P [56,57]. There are descriptions from other systems, where picophytoplankton biomass tended to increase even at high nutrient loading [58]. The observed dominance of small phytoplankton seems reasonable, even under elevated nutrient concentrations at the end of the experiment, as similar results were found in other shallow mesocosm experiments [59]. Furthermore, phytoplankton grew throughout the year in the shallow areas in proximity to the reed belt and possible P runoff of the DZLS, whereas there was only minor growth in deeper lagoon parts [11]. P stocks in the sediment were described to be highest in shallow areas [50], probably caused by large macrozoobenthos populations [60] who may increase nutrient burial through two- to three-fold increased sediment-water contact zones [61]. These results indicate that new production is generated in the shallow regions of eutrophic lagoons with high turn-over rates, instead of the whole turbid ecosystem. This gradient is not apparent through O_2_ production, but C accumulation. First, there was no significant difference of waterborne POC between both mesocosm treatments. Secondly, the share of water-borne POC compared to macrophyte-POC ranged from 22 – 67% at the end of the experiment (Supplement figure 2). These results indicated that phytoplankton production can be as high as macrophyte production in shallow areas. This mechanism can be missed, due to dilution, grazing on phytoplankton and macrophytes, or burial of biomass in the actual ecosystem.

Biomass built-up of macrophytes seems to be limited to patches in shallow margins of the ecosystem [11], as phytoplankton shading and invertebrate grazing [62] inhibits larger macrophyte biomasses. Sprouting in mesocosms took already two months at lower phytoplankton biomasses compared to the actual system and it can be inhibited if turbidity is too high [63]. Furthermore, macrophyte populations from phytoplankton-dominated systems seem to compensate high turbidity by faster sprouting at the extent of larger biomasses [64]. Therefore, only opportunistic, that means fast-growing, low-light adapted macrophytes will grow, if phytoplankton biomass is not depressed otherwise. This outcome will keep the system in a turbid state, as no macrophyte-mediated feedback mechanisms can develop.

## Conclusions

The focus of this study was to reveal and analyse production patterns in mesocosms representing shallow parts of a lagoon system, but without interference from higher trophic levels. Most strikingly, the depth-integration revealed a consistent net-heterotrophy, even in shallow, not-light limited mesocosms. Differences were observed due to different temperature acclimations of phytoplankton in shallow mesocosms compared to the actual ecosystem. This acclimation to changing environmental settings is probably a cause for the ongoing phytoplankton bloom even in the face of low nutrient availabilities. The occurring “boom-and-bust” cycles allowed a niche for annual, opportunistic submerged vegetation, as macrophytes proved to take over production gaps. However, our results indicate that phytoplankton biomass needs to be depressed further to allow proper macrophyte growth. Both, phytoplankton and macrophytes, seem to be of equal importance, when it comes to C-production over a season. This outcome highlights the capacity of shallow areas for ongoing eutrophication issues, as the shallow areas are the connection between the whole land-water transitional zone. Therefore, eutrophication issues may not be addressed by simple reduction of point sources, but restorations measures need to take nutrient burdens from the adjacent land into account. Furthermore, monitoring of coastal lagoon sites should include the waterfront, in combination with the water centre. Shallow areas seem to be “hot spots” in the spatial-temporal production pattern of eutrophic lagoon systems.

## Acknowledgements

We would like to thank the team of the Biological Station Zingst, namely R. Schumann, R. Wulff and V. Reiff for using the premise of the station, as well as analytical and set-up help. Furthermore, we would like to thank both working groups of ‘Marine Biology’ and ‘Aquatic Ecology’ of the University of Rostock for providing optodes. We thank H. Schubert, A. Schoor, and D. A. Campbell for discussions on that topic. This project was funded by the Federal Ministry of Education and Research. It was part of the project BACOSA I (grant: 03F0665A).

## Supplement

### Light and dark bottle experiment

On the 27^th^ August twelve bottles of 250 ml volume were filled in pairs from each mesocosm. One bottle was covered with aluminium foil and is referring to as the dark bottle, while the other one is called the light bottle. Both bottles were deployed into the same mesocosm were the water was taken from. The evolution of oxygen concentration was measured, after the initial oxygen levels were taken, (optode logger, Hq40, LDO, Hach-Lange) every two hours from 10:00 to 16:00 (CET). The dark bottles were kept in the mesocosm for 24h before measuring the final oxygen concentration. Percentages of oxygen saturation measured were converted in mg l^-1^ using solubility values (Benson and Krause 1984) and scaled to the bottles volume. Hourly rates of oxygen evolution were compared with hourly rates of oxygen evolution in mesocosm to cross-validate dissolved oxygen measurements. The results are shown in the supplement (Supplement. Fig S1).

Supplement Figure S1: Volumetric (μg O_2_ h^-1^) and Chlorophyll-normalized (μg O_2_ μg^-1^ Chlorophyll *a* h^-1^) net-community primary production (NCP) of plankton samples per Chlorophyll *a* concentration (μg l^-1^) within each mesocosm. Transparent and darkened bottles were used to measure oxygen concentration after every four hours and 24 hours. One pair of bottles is missing, as the plug of a dark-bottle loosened.

Supplement Figure S2: Particulate organic carbon (g m^-2^, based on 35 cm water depth) bound in either seston, or submerged macrophyte biomass on October 2014 (n = 3). Macrophyte mesocosms were left untouched until the end of the experiment. The POC value for macrophytes in phytoplankton mesocosms represents a cumulative sum of all previous harvests. Biomass in phytoplankton mesocosms was harvested four times from experiment start to end.

Supplement Table S1 Nutrient concentrations for dissolved inorganic phosphorus (DIP) and dissolved inorganic nitrogen (DIN, sum of ammonium, nitrate and nitrite) within mesocosms with and without macrophytes and the Zingster Strom from July to September. Standard deviation (±) was based on n = 3 in mesocosms and n=30 – 31 in the Zingster Strom.

Supplement Table S2 Characteristics of particulate material (seston) for Chlorophyll *a* (μg l^-1^), particulate organic carbon (POC, mg l^-1^), particulate organic nitrogen (PON, mg l^-1^), Carbon: Chlorophyll *a* (C/Chl *a*, mol C g Chl *a*^-1^), share of POC on total particulate matter (POC: seston in % based on mg l^-1^) in macrophyte mesocosms and phytoplankton mesocosms.

